# Lag Penalized Weighted Correlation for Time Series Clustering

**DOI:** 10.1101/292615

**Authors:** Thevaa Chandereng, Anthony Gitter

## Abstract

**Motivation:** The similarity or distance measure used for clustering can generate intuitive and interpretable clusters when it is tailored to the unique characteristics of the data. In time series datasets, measurements such as gene expression levels or protein phosphorylation intensities are collected sequentially over time, and the similarity score should capture this special temporal structure.

**Results:** We propose a clustering similarity measure called Lag Penalized Weighted Correlation (LPWC) to group pairs of time series that exhibit closely-related behaviors over time, even if the timing is not perfectly synchronized. LPWC aligns pairs of time series profiles to identify common temporal patterns. It down-weights aligned profiles based on the length of the temporal lags that are introduced. We demonstrate the advantages of LPWC versus existing time series and general clustering algorithms. In a simulated dataset based on the biologically-motivated impulse model, LPWC is the only method to recover the true clusters for almost all simulated genes. LPWC also identifies distinct temporal patterns in our yeast osmotic stress response and axolotl limb regeneration case studies.

**Availability:** The LPWC R package is available at https://github.com/gitter-lab/LPWC and CRAN under a MIT license.

**Supplementary information:** Supplementary files are available online.

## 1 INTRODUCTION

Time series data are extensively collected to study complex and dynamic biological systems [1, 2]. Tracking the levels of biological molecules such as genes and proteins over time can reveal interactions among them [1] and inform treatment decisions in various diseases [3]. Temporal or longitudinal data are important across multiple disciplines (for example, finance, engineering, and medicine), but biological time series datasets are often shorter than those in other domains. Typically, separate experiments are required for each time-point, which limits the number of timepoints collected.

Similarity in gene expression patterns can correspond to similarity in biological function, which helps direct future research [4]. Countless clustering algorithms group data points with similar characteristics, but the meaning of “similar” is inherently subjective and application-specific. In time series datasets, similarity must account for the temporal structure. Unlike other data types, observations in time series datasets are dependent on the past. General purpose clustering methods may be able to detect synchronized temporal changes over time but cannot recognize that two entities have the same temporal profile if one is delayed or lagged after the other. In addition, in many cases the time-points in a biological study are not uniformly distributed over time, and the selection of timepoints is an important aspect of the experimental design [5]. The duration between timepoints in irregular time series affects the similarity of temporal profiles, especially when allowing lags among the clustered entities.

Many time series clustering algorithms have been introduced to understand the dynamics of biological processes. Clustering methods can be divided into two branches: hierarchical and partitioning [6]. Hierarchical clustering methods either build small clusters that iteratively merge into larger clusters or large clusters that divide into smaller clusters [1]. Partitioning, on the other hand, divides entities according to their specific characteristics and often requires specifying the number of clusters in advance.

Hierarchical clustering methods, such as clustering with correlation or transformed Euclidean distance for similarity, were a common choice before the proliferation of time series-specific algorithms [4] and continue to be widely used for temporal data [7]. Generic approaches ignore the sequential nature of the data and give the same clusters even if the timepoints are shuffled, but many temporal hierarchical clustering methods exist. Dynamic Time Warping (DTW) is designed for biological time series data where the data is locally aligned so that the Euclidean distance is minimized [8, 9]. Aach et al. introduced simple algorithms for time warping that interpolate intermediate intensities between time-points and align them accordingly [9]. Short time series (STS) distance accounts for the sequential nature of temporal data by computing the rate of change in intensity between adjacent timepoints, but it does not consider lags [10]. TimeClust implements two clustering algorithms, Temporal Abstraction Clustering (TAC) and Random Walk Models for Bayesian Clustering, developed specifically for short time series [11, 12, 13]. Neither of them accounts for lags.

Many partition-based clustering algorithms are available for biological time series data. The Short Time-series Expression Miner (STEM) enumerates temporal template profiles and matches genes to them so it works best for short time series (3-8 timepoints) [14]. DynaMiteC [15] clusters genes by fitting them to prototype impulse models [16], but impulses are only one type of common temporal pattern [1]. DynOmics uses fast Fourier transform to model expression values using mixtures of cyclic patterns [17]. This method also realigns expression values to account for delays but does not treat lagged and unlagged genes differently. Graphical Query Language (GQL) clusters based on a hidden Markov model [18]. Bar-Joseph et al. turn discrete time series expression data into continuous data using splines [19]. Their clustering algorithm uses the continuous data and expectation maximization to optimize alignment of the temporal data. Other partitioning-based algorithms include a wavelet-based density method using multi-level thresholding [6] and Cluster Analysis of Gene Expression Dynamics (CAGED), which uses autoregressive equations [20].

Other time series clustering methods are Bayesian models [21, 22]. Several of these are built on Dirichlet processes with mixture models that use the temporal information [7, 23]. Dahl proposes a clustering algorithm where genes with similar Dirichlet process mixture components are grouped together and the model is fit using Markov Chain Monte Carlo [23]. McDowell et al. use a Dirichlet process Gaussian process mixture model as the posterior distribution and a Gaussian process as the prior to cluster time series data [7].

Despite the abundance of clustering algorithms, popular clustering methods do not capture important temporal properties such as lags and irregular timepoints, which we demonstrate with a simple toy example. Figure 1 shows how four artificial gene expression profiles are grouped by different clustering methods. Hierarchical clustering with Euclidean distance (heuc) ignores the timing of the spikes entirely. Existing time series clustering algorithms also fail to group the early and late genes. STS captures the difference in intensity with the prior timepoint, and DTW optimizes the distances using time shifts. We introduce a time series clustering algorithm, Lag Penalized Weighted Correlation (LPWC), which captures the delayed responses and the similarity of the early and late genes. LPWC has two modes with a high lag penalty (hLPWC) and low lag penalty (lLPWC).

One of the main contributions of LPWC is a similarity function that accounts for pairs of temporal profiles that may occur at slightly different times. This generates a gene-gene similarity matrix that can be used as input for standard similarity- or distance-based clustering methods such as hierarchical clustering. The LPWC similarity score is derived from weighted correlation, but the correlations of lagged temporal profiles are penalized using a Gaussian kernel. The kernel is also used to account for irregular time sampling. We demonstrate the advantages of LPWC over existing general and time series clustering algorithms on a simulated impulse model dataset and case studies on the yeast osmotic stress response and axolotl limb regeneration.

**Figure 1.**
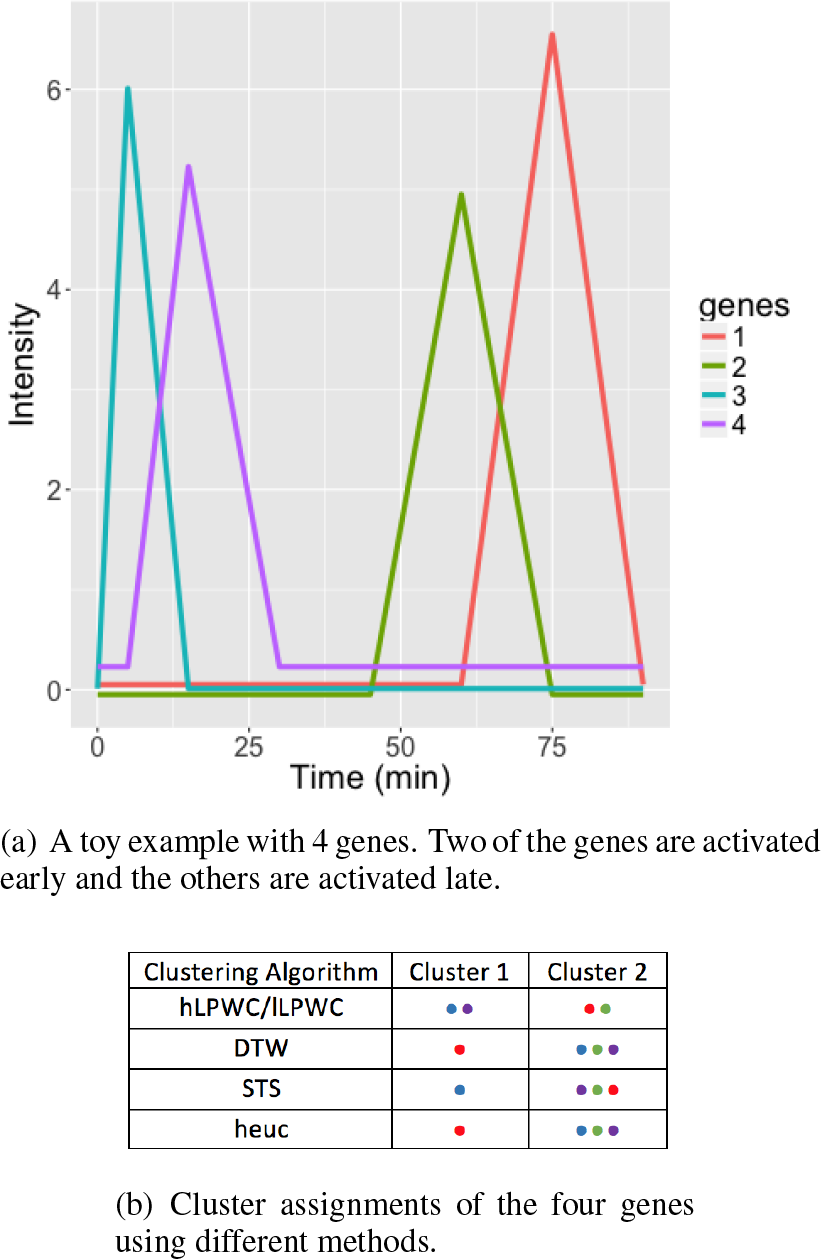
A toy example with four genes and timepoints at 0, 5, 15, 30, 45, 60, and 75 mins (Figure 1(a)). Each of the genes has a sharp rise and fall in expression, which occurs at a different time-point. Genes 1 and 2 both have late spikes and intuitively should be clustered together. Genes 3 and 4 are both early. Several widely used clustering methods group the genes into two clusters (Figure 1(b)), but only LPWC groups the early and late genes. The colored dots represent the different genes in Figure 1(a).

## 2 METHODS

### 2.1 Lag Penalized Weighted Correlation

The goal of LPWC is to group genes that have similar shapes in their expression levels over time. These shapes or temporal profiles refer to the patterns of increases and decreases in expression. Two genes have similar temporal shapes if the timing of these increases and decreases coincides even if the expression levels are not the same. In order to identify similar temporal shapes that are not perfectly synchronized, LPWC applies a lag operator to re-align the timepoints when comparing two expression profiles (Figure 2(a)). The lag operator compares the timepoints of one expression profile with later timepoints in the other profile. LPWC only considers the expression levels at the observed timepoints and does not interpolate between timepoints or rescale time as in DyNB [24]. Interpolation with line segments [9] or splines [19, 25, 26] makes assumptions about the unobserved behavior between timepoints. Gaussian processes make much weaker assumptions [7], but the kernel function still constrains which types of temporal behaviors and smooth profiles are most likely in between the observed times [27]. Because the aligned time series can pair measurements that are temporally far apart, LPWC weights the pairs of timepoints to give stronger consideration to those that are close in time.

**Figure 2.**
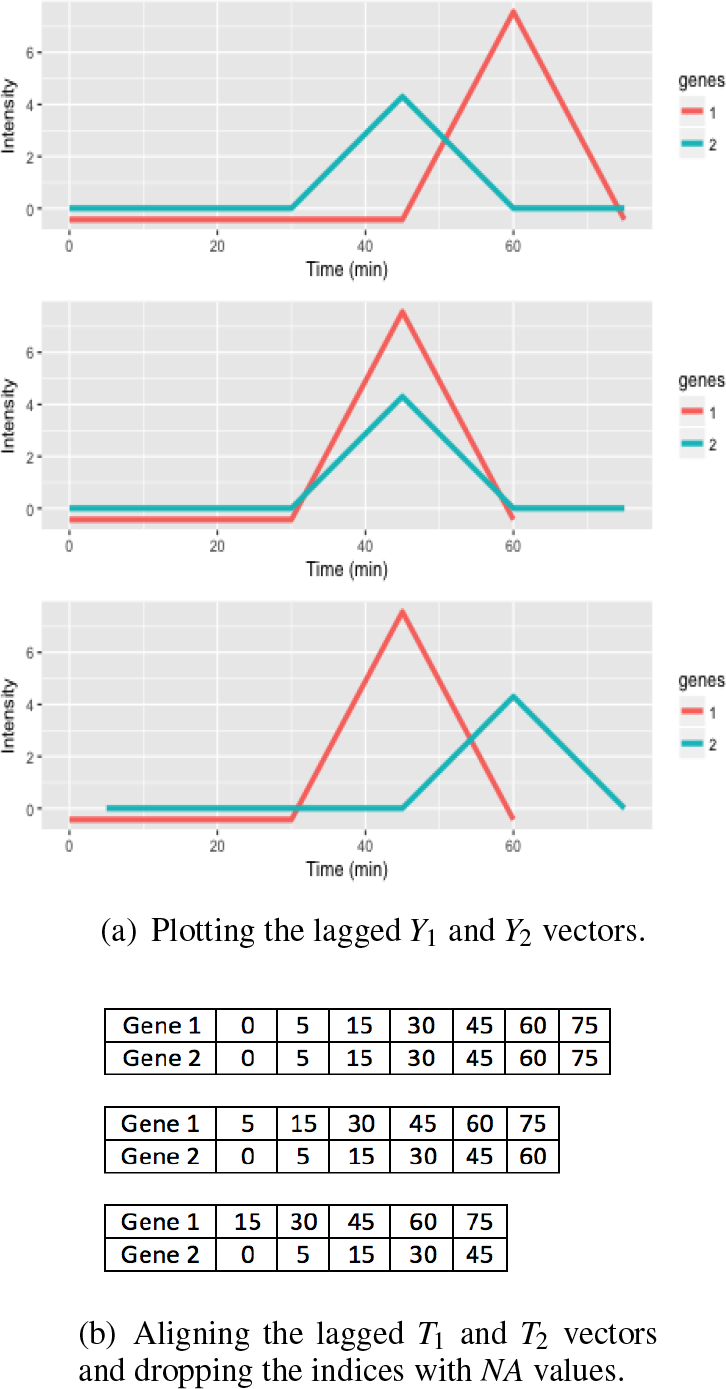
An example of the effects of applying different lags to genes 1 and 2. Figure 2(a) shows aligned expression vectors and Figure 2(b) shows the aligned timepoints. With no lags, *X*_1_ = 0 and *X*_2_ = 0, the temporal profiles of genes 1 and 2 are not aligned so the gene pair will have a low LPWC similarity score (top row). With lags *X*_1_ = −1 and *X*_2_ = 0, the patterns are aligned, and the LPWC similarity score will be high (middle row). Finally, with *X*_1_ = −1 and *X*_2_ = 1, the temporal shapes are once again not aligned, and the LPWC similarity score will be even lower than in the no lag case because the penalty for introducing lags is applied (bottom row).

The LPWC algorithm is composed of three steps: choosing lags for each gene, computing the final similarity matrix for all gene pairs, and running standard hierarchical clustering. The best lags are selected by maximizing the sum of the similarities for each gene with respect to all other genes. The maximum possible gene-gene similarity decreases as the lag in the aligned timepoints increases because we prefer to recover synchronized temporal behaviors or similar temporal patterns that are close in time. In addition, we have less confidence in the similarity of the temporal shapes when they are computed with shorter temporal subsequences. The final correlation-based similarity is computed once all the lags are fixed.

The overall correlation-based similarity function of each gene pair *i*, *j* is computed as below.

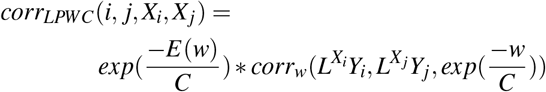

where *L* is a lag operator, *X_i_* is the lag for gene *i*, *Y_i_* is the temporal expression levels of gene *i*, *C* is a parameter that controls the lag penalty, *w* is a weight vector, and *corr_w_* is a weighted correlation. The lag *X_i_* is an integer-valued variable that represents the number of indices a temporal profile is shifted forward or backward in time, where positive values represent forward shifts. The lag operator *L* can be applied to a vector of temporal gene expression levels (*Y_i_*) or a vector of timepoints, which we denote as *T_i_* for gene *i*. *L* reduces the effective length of the lagged vector, introducing *NA* placeholder values. For example, if *T_i_* = [0,5,15] and *Y_i_* = [0.2,1.4,4.5], then for *X_i_* = 1 we have *L^1^T_i_* = [*NA*, 0,5] and *L^1^Y_i_* = [*NA*, 0.2,1.4]. For *X_i_* = −1 we obtain *L*^−1^ *T_i_* = [5,15,*NA*] and *L*^−1^*Y_i_* = [1.4,4.5,*NA*].

Given this lag operator, we can define the weight vector w for weighted correlation.

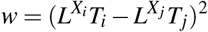

The vector subtraction is performed after dropping indices where either vector is *NA*. Similarly, in the weighted correlation, defined here generically for input vectors *x* and *y* and weight vector *z*,

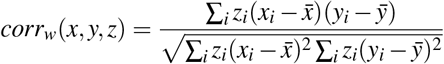

we drop indices that are *NA* in either *x* or *y* (Figure 2(b)).

The overall penalty for aligning timepoints is derived from the mean weight *E*(*w*).

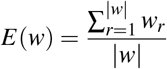

where *r* is an index for the elements of the weight vector and |*w*| is the vector length. When *X_i_* = 0 and *X_j_* = 0, *w* is the zero vector and *E*(*w*) = 0. Thus, the special case *corr_LPWC_*(*i*, *j*, 0, 0) is the standard (unweighted) Pearson correlation of *Y_i_* and *Y_j_*.

Because choosing the optimal lags *X_i_* for all genes is NP-complete (see Supplementary Section 4.1 for a sketch of the proof), we use a heuristic approach. For each gene *i*, we store the score and respective lag with respect to all other genes *j*. The parameter *m* is the maximum lag allowed in the data. It is important to control the maximum lag because lags reduce the number of data points used to calculate the weighted correlation.

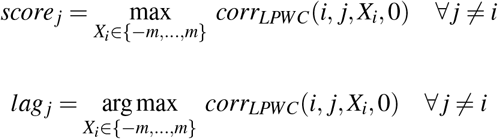

Then, a best lag 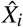 for gene *i* assigned by

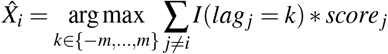

where *I* is an indicator function. This is repeated to select a best lag for all genes.

Upon obtaining the best lags 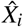 for all genes, we compute the similarity

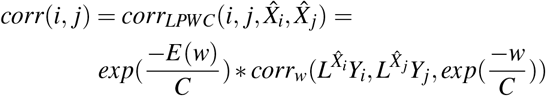

The similarity measure *corr*(*i*, *j*) can be used directly by a clustering algorithm that requires gene-gene similarities as input. However, LPWC uses hierarchical clustering, which requires a distance measure instead. We know that −1 ≤ *corr*(*i*, *j*) ≤ 1. Thus, we transform the similarities with *dist*(*i*, *j*) = 1 − *corr*(*i*, *j*) to obtain distances for hierarchical clustering.

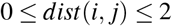

We run hierarchical clustering with complete linkage.

### 2.2 Controlling the lag penalty

Because lags reduce the number of timepoints used for the correlation calculation and biological time series data are typically short already, there is a risk that two lagged expression vectors will have a high correlation score by chance. Lagged correlation clustering without modification does not perform well [17]. Thus, a Gaussian kernel 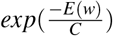 is used to scale and penalize the weighted correlation based on the lags. The parameter *C* controls the width of the Gaussian kernel function and the severity of the penalty. The appropriate *C* is subjective and application-specific. Instead of choosing one universal default penalty parameter *C*, LPWC implements two data-dependent ways to set *C*: the high and low penalty modes. The high penalty (hLPWC) penalizes lags more, increasing the possibility of setting *X_i_* = 0 compared to the low penalty (lLPWC), which will set more *X_i_* ≠ 0. In addition to these two default options, the user can also specify *C* directly to introduce more or fewer lags.

The overall penalty that LPWC applies to the weighted correlation *corr_w_* is 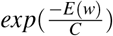, which scales the correlation by a factor between 0 and 1. For the high penalty, we set the mean penalty over all valid positive lags to 0.5 and solve for *C*

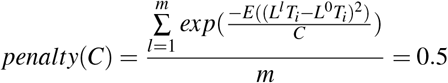

where *m* is the maximum lag and (*L^l^T_i_* − *L*^0^*T_i_*)^2^ is the weight vector *w* when comparing the timepoint vector *T_i_* with a lagged version of *T_i_*.

For the low penalty, we compute the values of *C* for which *penalty*(*C*) produces penalties between 0.5 and 0.95 with a step size of 0.05. For each of those *C*, we run LPWC and obtain the gene-gene similarity matrix. We choose the *C* for which the gene-gene similarity matrix is the most stable with respect to the similarity matrix from the previous *C*. Stability is computed by subtracting the two gene-gene similarity matrices, squaring the elements, and summing them. The lowest sum squared error is preferred. Because it sweeps over multiple values of *C*, lLPWC is slower than hLPWC.

### 2.3 Comparison with existing methods

The clustering algorithms used for comparison are Euclidean distance with hierarchical clustering (heuc) and kmeans clustering (keuc), dynamic time warping with hierarchical clustering (DTW), short time series distance with hierarchical clustering (STS), and Pearson correlation with hierarchical clustering (hcorr) and kmeans clustering (kcorr) (Supplementary Section 4.2). These algorithms include some of the most widely used general approaches as well as two tailored for time series.

### 2.4 Simulated time series

To test LPWC, we simulated time series gene expression data using an impulse model called ImpulseDE [16]. Impulses are one common type of temporal pattern in gene expression data [1]. An impulse can be represented as a parameterized curve in which each gene has an initial expression level, increases or decreases in response to a hypothetical stimulus, and then rises or falls to a new steady state level. The impulse model parameters control each level, the timing of the expression increases and decreases, and the curvature of the expression changes (Table S1).

### 2.5 Cluster evaluation

Cluster evaluation is difficult because in general the true clusters are not known. The rand index has been used traditionally to evaluate similarity between two clustering results [28]. However, to control for randomness and compare clustering scores from clusters of different sizes, adjusted rand index (ARI) is a more suitable metric [28]. ARI yields a score of 1 for a perfect clustering that matches the true cluster labels. On the other hand, a score close to 0 indicates a poor clustering.

**Figure 3.**
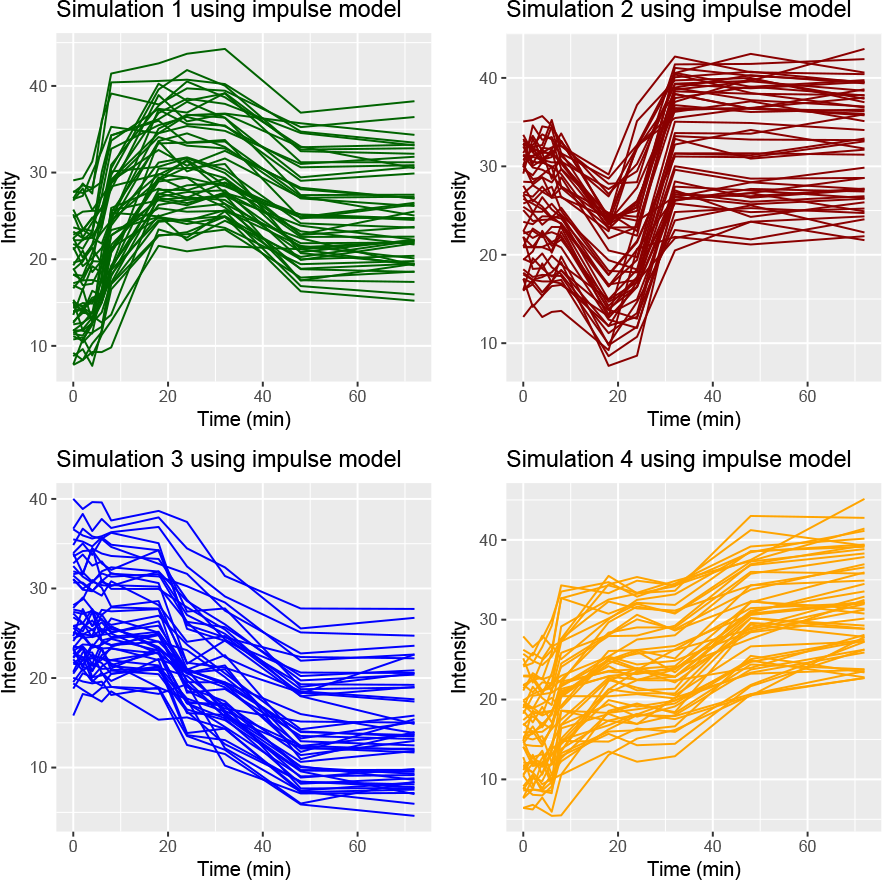
Four models simulated using ImpulseDE. Each model has different characteristics (expression increases and decreases over time) and contains 50 genes. The initial ImpulseDE model parameters were randomly modified when sampling each gene to induce variation. In addition to randomly modifying the parameters, random noise sampled from the Gaussian distribution N(0, 1) was added to the simulated expression profiles.

One way to evaluate time series clustering algorithms is by assessing how important the temporal information is to the clustering results. We obtain clusters using the original data and then permute the data by randomly reordering the timepoints (the gene expression observations do not change). The permutations destroy the true temporal dependencies in the data. If a clustering algorithm does not use the temporal information, the ARI score when comparing its clusters on the original and permuted data will be close to 1, which is undesirable. We repeat the timepoint permutation 100 times for each clustering algorithm and assess the distribution of ARI scores.

Another challenge is choosing the number of clusters, which can be addressed with the silhouette method [29]. This method assesses whether the clusters are cohesive and distinct from one another. We select the number of clusters that maximizes the average silhouette width.

### 2.6 Case studies

We applied LPWC in two case studies to demonstrate how it can be used to obtain coherent temporal clusters and derive biological insights into dynamic transcriptional and signaling processes. The first captures the rapid phosphorylation response to osmotic stress in yeast [30]. Kanshin et al. obtained mass spectrometry-based phosphorylation samples in NaCl-induced osmotic stress and control conditions, uniformly sampling 0 to 60 seconds post-stimulation every 5 seconds for a total of 13 timepoints. They transformed these into log2 stress versus control fold changes at each timepoint. We clustered the 344 singly phosphorylated phosphopeptides that were reported to have significant dynamic changes and were not missing values at any timepoints.

The second dataset contains time course RNA-seq data from the axolotl blastema after amputating the right forelimb [31]. Stewart et al. studied the transcriptional changes that take place during the transitions from wound healing to dedifferentiation to limb regeneration. They sampled gene expression at 12 timepoints: 0, 3, 6, and 12 hr and 1, 3, 5, 7, 10, 14, 21, and 28 days post-amputation. Unlike the osmotic stress application, there is drastic irregularity between consecutive sampling times. We converted all times to days. Because the axolotl genome had not been sequenced at the time, Stewart et al. mapped axolotl contigs to human transcripts. They processed the data using edgeR [32], comparing each timepoint to the 0 day measurement to obtain the up- and down-regulated genes. A total of 1656 genes were up- or down-regulated at least at one timepoint compared to 0 day. We ran LPWC on their mapped human gene expression data.

### 2.7 Gene enrichment analysis

We performed gene enrichment analysis of the LPWC cluster members in DAVID 6.8 [33, 34] (Supplementary Section 4.3). For the yeast and axolotl case studies, we report Gene Ontology (GO) GOTERM_BP_FAT [35] and KEGG-PATHWAY [36] terms that are enriched using DAVID parameters Counts = 2 and Ease = 0.05. The terms were further filtered for false discovery rate ≤ 5%.

### 2.8 Software availability

LPWC is implemented in R and released as open source software under the MIT license. The LPWC package is available at https://github.com/gitter-lab/LPWC and CRAN. We used LPWC version 0.99.0 for all analyses.

## 3 RESULTS

To assess LPWC, we compared it to other popular clustering algorithms on a simulated time series dataset where the true clusters are known and conducted two biological case studies. The yeast osmotic stress response data consist of NaCl induced osmotic stress phosphorylation samples obtained from mass spectrometry [30]. The axolotl blastema RNA-seq data is collected upon amputating the right forelimb [31].

### 3.1 Simulated time series data

The simulated time series data contains 200 genes with 10 timepoints: 0, 2, 4, 6, 8, 18, 24, 32, 48, and 72 mins. The data is composed of four distinct temporal patterns with 50 genes per pattern. Because the true pattern used to generate each temporal profile is known, the ARI score can be obtained by comparing the true clusters to the cluster assignments produced by different clustering algorithms.

We evaluate four general clustering algorithms (heuc, keuc, hcorr, and kcorr), two time series clustering algorithms (DTW and STS), and LPWC. Instead of using the silhouette method to pick the number of clusters, all methods return exactly four clusters, the correct number of clusters from the simulation. The two LPWC variants with high and low lag penalties, hLPWC and lLPWC, outperform all other methods (Figure 4). The clusters from hLPWC (Figure S1) and lLPWC (Figure S2) show that the simulated genes are accurately clustered according to the known assignments in Figure 3. The ARI scores obtained are close to 1, a nearly perfect clustering. However, the time series clustering methods DTW and STS perform poorly on this task, and hcorr and kcorr are the only other methods that perform reasonably well. This simulation acts as a positive control, demonstrating that LPWC correctly recovers the four temporal expression patterns when we insert offsets in the timing and expression levels.

**Figure 4.**
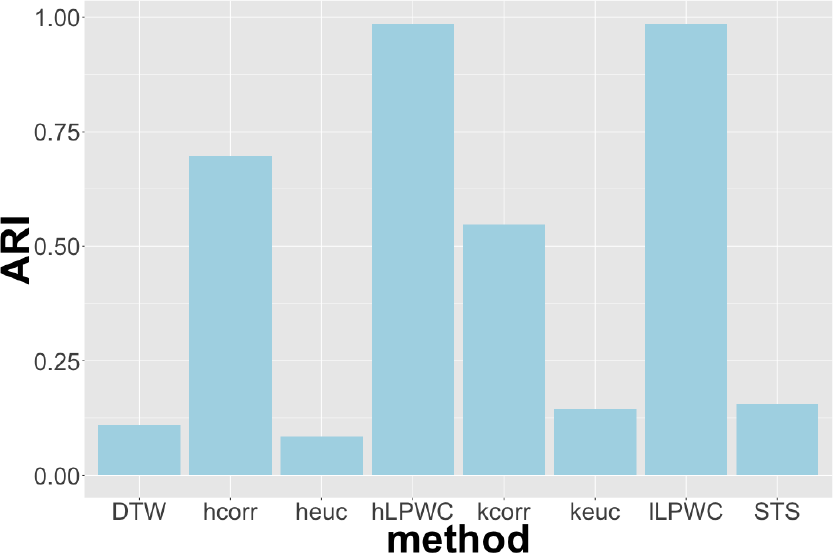
ARI scores for different clustering methods for the simulated data where the real clusters are known.

### 3.2 Case studies

For biological case studies, the true clusters are not known, and it is harder to quantitatively evaluate clustering methods. Therefore, we assess whether each clustering algorithm produces clusters with discernible common temporal patterns and makes use of the temporal structure in the data. To assess how the temporal structure is used during clustering, we permute the timepoints.

#### 3.2.1 Yeast osmotic stress response

We used lLPWC to cluster the yeast phosphopeptides in the osmotic stress response dataset into three clusters (Figure 5 and Supplementary File 1). Although cluster 3 contains fewer phosphopeptides than the others, this number of clusters was optimal based on our silhouette analysis in Figure S3. Clusters 1 and 2 were comparable for both lLPWC and hLPWC, with only the smaller cluster 3 showing a notable difference in the mean temporal trend (Figures S4 and S5 and Supplementary File 2).

There are 33 nonzero lags in lLPWC (Table S2) among the 344 phosphopeptides and 26 nonzero lags in hLPWC (Table S3). Although there are not many lags introduced, they are important in aligning the temporal structure of the phospho-peptides with other phosphopeptides. All clusters exhibit distinct temporal patterns. For visualization purposes, we emphasize these patterns by subtracting the value at 0s from all timepoints before applying the lags and plotting the cluster members and the mean temporal trend (Figures 5 and S5). In both lLPWC and hLPWC, most of the phosphopeptides are assigned to clusters 1 and 2 (Tables S4 and S5). Phosphopeptides in cluster 1 demonstrate an overall increasing trend over time. The cluster members are enriched for many broad GO terms related to signal transduction, cellular response to osmotic stress, and actin cytoskeleton organization, which was previously reported to be an important component of this stress response [30] (Supplementary File 1). Cluster 1 includes the mitogen-activated protein kinase Hog1 and other important proteins in the osmotic stress response pathway such as kinases Pbs2 and Rck2 and transcription factors Msn4 and Sko1. Cluster 2 phosphopeptides show a decrease in phosphorylation over time. The steady increase and decrease trends in clusters 1 and 2 also reflect the major patterns reported by Kanshin et al. [30]. Similar to cluster 1, cluster 2 is also enriched for actin-related terms and general signaling as well as salt and osmotic stress response proteins. These include additional transcription factors Cin5 and Msn2 as well as different phosphorylation sites on Msn4 and Pbs2. Cluster 3, though small, contains a group of phosphopeptides with mostly small changes in phosphorylation over time except for a distinct decrease and increase at 45s. This cluster contains cytokinesis-related proteins.

**Figure 5.**
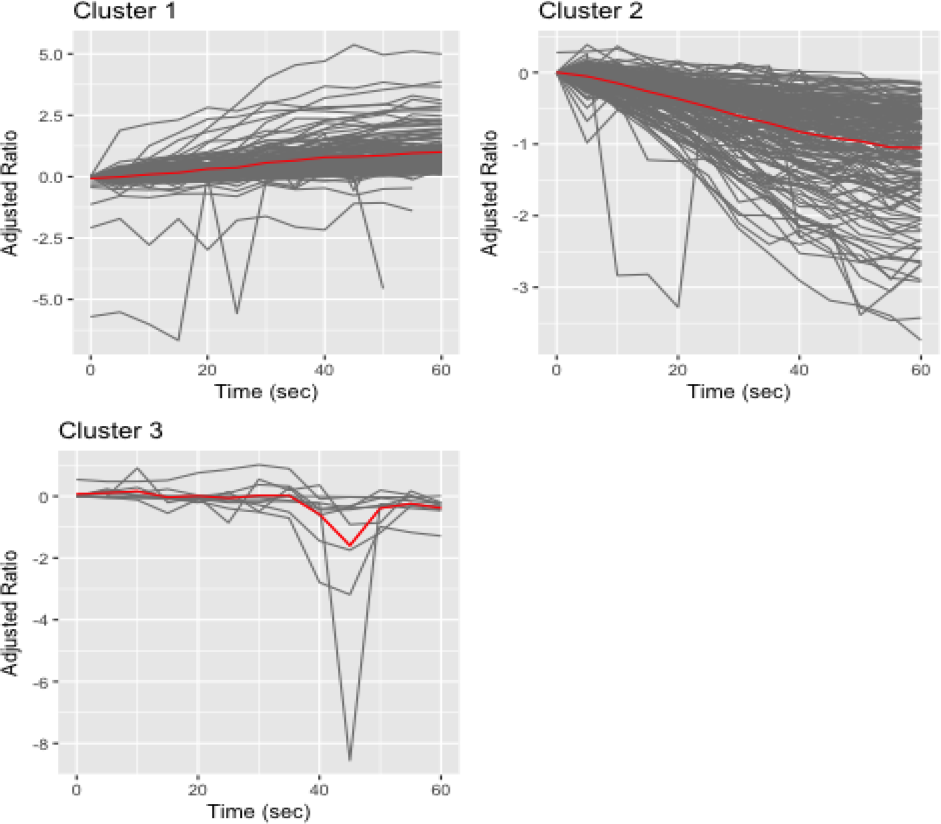
Clusters for the yeast data using the lLPWC algorithm. The y-axis shows the log2 salt/control ratio after subtracting the 0s log2 ratio from all values so all temporal profiles start at 0.

When comparing the real and permuted data (Figure S6 and Table S6), nearly all of the hlPWC and lLPWC ARI scores are greater than 0.75, which reflects the relatively small number of lagged genes in the real data. As expected, the four general clustering algorithms (heuc, hcorr, kcorr, and keuc) do not use the temporal information and have ARI scores of 1. STS has a low ARI but performs poorly on this dataset. It places all genes into a single cluster, except for two genes that are each assigned to their own singleton cluster (Figures S7 and S8 and Table S7). DTW does quite well on the yeast data (Figures S9 and S10). Although three of its clusters are small (Table S8), the other five contain distinct temporal patterns. The ARI score is also low, showing that DTW does account for the temporal structure in the yeast osmotic stress response data.

#### 3.2.2 Axolotl blastema

Figure 6 shows the time series plot of the axolotl blastema gene expression clusters from our hLPWC algorithm. Three clusters were selected based on the average silhouette width in Figure S11, and the number of lagged genes and cluster sizes are reported in Table S9 and Table S10, respectively. We added 1 to each expression value before taking the log2 ratios with respect to time 0 days for visualization purposes only. This dampens the extreme fold changes that occur when the initial gene expression level is close to 0, which obscure the temporal trends in each cluster.

The major temporal trends in each cluster are similar for hLPWC (Figure 6 and Supplementary File 3) and lLPWC (Figures S12 and S13, Tables S11 and S12, and Supplementary File 4). In cluster 1, gene expression rapidly then gradually increases from 0 to 14 days, decreases until the 21 day timepoint, and stabilizes afterward. This cluster is enriched for GO terms related to the cell cycle, proliferation, blood vessel development, and wound healing (Supplementary File 3). The mean cluster 2 trend exhibits down-regulation from 0.25 days until 14th days, and the cluster is associated with GO and KEGG terms involving ribosomes, RNA-related metabolism, muscle development, and response to oxidative stress. In cluster 3, the mean expression decreases immediately and then rises until day 1, at which point it decreases again until day 10 and then increases for the remainder of the duration. These genes are enriched for type I interferon signaling and other immune processes.

Because hundreds of genes are assigned non-zero lags in the axolotl case study, both hLPWC and lLPWC have low ARI scores when comparing their clusters with the permuted data (Figure S14 and Table S13). As with the yeast case study, the four general clustering methods have ARI scores of 1 or close to 1. STS performs poorly once again. One cluster contains 99% of the genes despite their different temporal characteristics (Figure S15 and S16 and Table S14). Although DTW was excellent on the yeast dataset, it struggles with the axolotl data (Figure S17 and S18 and Table S15). The ARI scores are much higher than on yeast, and it places 98% of genes in a single cluster.

**Figure 6.**
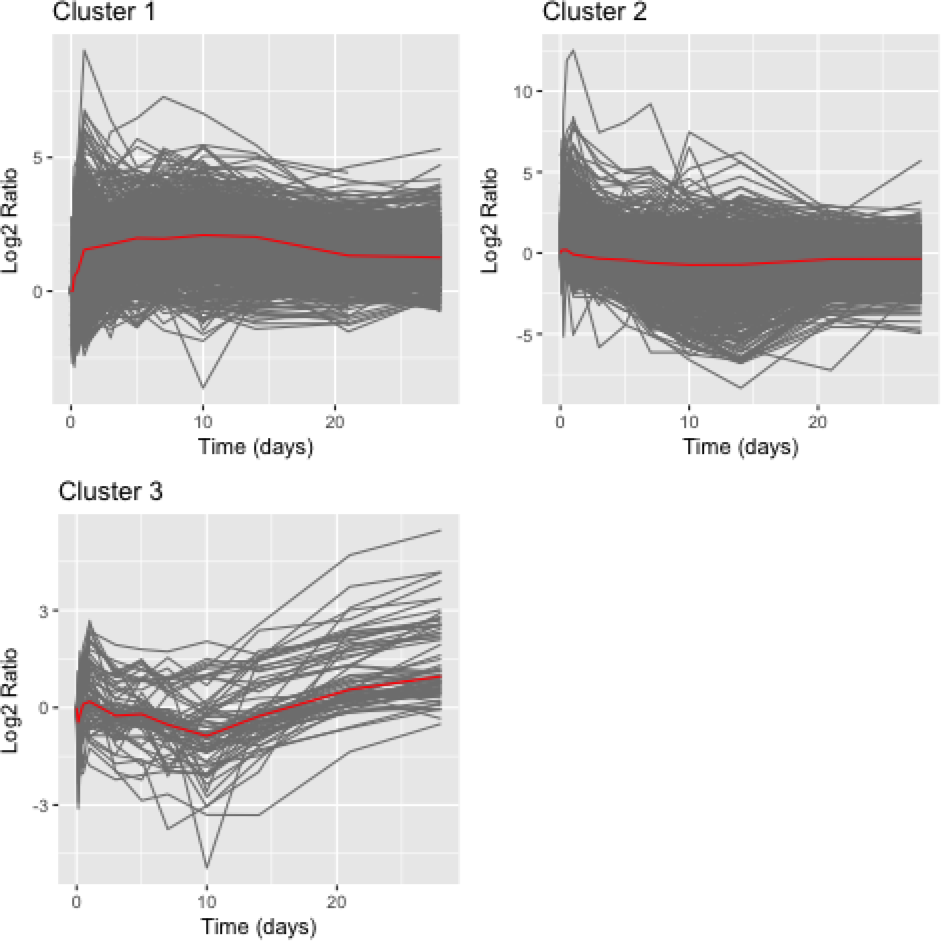
Clusters for the axolotl data using the hLPWC algorithm. The log2 ratio is with respect to the 0 day timepoint.

## 4 DISCUSSION

LPWC is designed to capture temporal structure when clustering biological datasets, specializing in irregularly-sampled time points and detecting delayed responses. It uses lags to align temporal gene expression profiles and weighted correlation to account for irregular sampling. The similarity scores of lagged genes are penalized in order to prefer synchronized temporal patterns and correlations that are computed using a greater fraction of the timepoints. The hLPWC option captures fewer lags than lLPWC due to the higher penalty that is imposed. The choice of lags is important because the observations at the beginning and end of the time series are dropped when comparing aligned lagged genes. For instance, in Figure 1(a), assigning gene 4 a lag of -1 gives it the same temporal pattern as gene 3 but ignores that gene 4 increases in expression before decreasing. Therefore, the default parameters are conservative, which is why only a small fractions of phosphopeptides or genes are lagged in our yeast case study.

In both the yeast and axolotl case studies, LPWC successfully identifies clusters with unique temporal patterns. LPWC introduces more lags in the axolotl dataset, which may be due to the close timing of the initial timepoints, and the temporal permutation analysis reflects the stronger temporal dependency when there are more lags. The general kmeans and hierarchical clustering algorithms disregard the timepoints by design. STS has performed well in other applications, but places nearly all phosphopeptides or genes into a single cluster in both case studies here. DTW on the other hand does very well on the yeast application, but like STS creates a single dominant cluster on the axolotl dataset. Only LPWC produces useful clusters that depend on the timing for both of the datasets. In addition, it is the only method that accurately recovers the correct clusters in our simulated data.

Because the main advantage of LPWC is its ability to introduce lags to detect common but unsynchronized temporal patterns, it works best when there are sufficient timepoints to support multiple lags. At least four timepoints are required by LPWC in order to allow one lag. However, if gene *i* and gene *j* are assigned lags of -1 and 1, only two timepoints remain to estimate the correlation. Thus, it is advisable to use LPWC with five or more timepoints. STEM [14], which enumerates temporal patterns, may be preferable for very short time series datasets without delayed responses. DTW [8, 9] has been used extensively in the financial industry for long time series datasets. LPWC can also be applied to long time series, but the default lag penalties will prevent large lags from being introduced.

LPWC is a generalization of traditional hierarchical clustering with correlation-based similarity. In the case where no lags are introduced, LPWC is identical to standard hierarchical clustering. With the default lag penalty settings, LPWC can estimate how many genes should be lagged based on the characteristics of each particular dataset. Various clustering algorithms have been built upon distance-based or correlation-based similarity measures. These approaches capture different types of temporal shapes. Correlation reveals the trends in the data whereas distance captures the difference in magnitude of expression levels or fold changes. The preference for one over the other is subjective. We prefer correlation for LPWC because it can be applied directly to the original expression levels without computing fold changes with respect to the initial timepoint. Distance-based time series clustering often requires computing these fold changes so that genes are grouped based on their temporal patterns instead of their average expression level, but this effectively drops one of the timepoints because the variation at the initial timepoint is ignored.

Currently, LPWC only accepts a single dataset with one set of timepoints. One future direction would be to support clustering mixed datasets with different timepoints, which could be especially useful when combining multiple types of biological data. There are also opportunities to better approximate the NP-complete lag optimization problem (Supplementary Section 4.1) and estimate the default value of C, which controls how many lags are introduced.

## 5 ACKNOWLEDGEMENTS

This research was supported by NSF CAREER award DBI 1553206, the University of Wisconsin Carbone Cancer Center Support Grant NIH P30 CA014520, and the UW-Madison Center for High Throughput Computing in the Department of Computer Sciences. We are grateful to Ron Stewart, James Dowell, Karl Broman, Wenzhi Cao, Jen Birstler, Atul Deshpande, Shengchao Liu, and all members of the Gitter lab for their helpful feedback and discussions.

